# Convergence in sympatry: evolution of blue-banded wing pattern in *Morpho* butterflies

**DOI:** 10.1101/2020.05.13.093096

**Authors:** V Llaurens, Y Le Poul, A Puissant, C Noûs, P Blandin, V Debat

**Author notes:** Corresponding author: Violaine Llaurens.

## Abstract

Species interactions such as mimicry can promote trait convergence but disentangling this effect from those of shared ecology, evolutionary history and niche conservatism is often challenging. Here by focusing on wing color pattern variation within and between three butterfly species living in sympatry in a large proportion of their range, we tested the effect of species interactions on trait diversification. These butterflies display a conspicuous iridescent blue coloration on the dorsal side of their wings and a cryptic brownish colour on the ventral side. Combined with an erratic and fast flight, these color patterns increase the difficulty of capture by predators and contribute to the high escape abilities of these butterflies. We hypothesize that, beyond their direct contribution to predator escape, these wing patterns can be used as signals of escape abilities by predators, resulting in positive frequency-dependent selection favouring convergence in wing pattern in sympatry. To test this hypothesis, we quantified dorsal wing pattern variations of 723 butterflies from the three species sampled throughout their distribution, including sympatric and allopatric situations and compared the phenotypic distances between species, sex and localities. We detected a significant effect of localities on colour pattern, and higher inter-specific resemblance in sympatry as compared to allopatry, consistent with the hypothesis of local convergence of wing patterns. Our results provide some support to the existence of escape mimicry in the wild and stress the importance of estimating trait variation within species to understand trait variation between species, and to a larger extent, trait diversification at the macro-evolutionary scale.

## Introduction

Understanding the evolutionary forces driving trait diversification between species is challenging, as it results from a combination of contingent historical events, neutral divergence and adaptive evolution. Disentangling the effect of neutral divergence from the effect of selective forces that could be either shared or contrasted between species is a major question for evolutionary biologists. Selective forces shared by closely-related species living in similar ecological niches can limit divergence in adaptive traits, resulting in niche conservatism along phylogenies (Wiens *et al*. 2010). On the contrary, when closely-related species live in sympatry, trait divergence can be promoted by selection caused by species interactions, including competition for resources (Schluter 2000) or reproductive interference (Gröning & Hochkirch 2008). Documenting the relative importance of niche conservatism among closely-related species *vs*. character displacement is essential to comprehend trait diversification between species, and to estimate how much species interactions shape macro-evolutionary patterns of trait variation.

Closely-related species partly living in sympatry offer a great opportunity to disentangle the effects of species interactions from those of shared selective pressures acting on the evolution of their traits: in geographic areas where several closely-related species live in sympatry, the evolution of traits within a species can be influenced by the evolution of traits in sympatric species, while the selective forces generated by species interaction are no longer acting in allopatry. The role of species interactions in trait divergence in sympatry has been well-documented for mating cues and preferences, as for instance in the flycatcher *Fiducela hypoleuca* where plumage coloration is divergent from the ancestral dark coloration in a population where the dark sister-species *F. albicolis* lives in sympatry, as the result of selection against hybrids (Strae *et al*. 1997). Other antagonistic interactions such as resources competition have also been reported to drive character displacement in sympatry, as illustrated by the change in beak size in island population of the Darwin finch *Geospiza fortis* following the arrival of the competitor species *G. magnirostris* (Grant & Grant 2006). While the effects of antagonistic interactions on trait divergence in sympatric species have been well-documented, those of mutualistic interactions remain scarcely studied. Evidences from a few obligate mutualisms such as fig-waps pollination (Jousselin *et al*. 2003) or acacia-ants protection (Ward & Branstetter 2017) have nevertheless highlighted that positive interactions can drive repeated evolution in traits involved in the interaction, such as ostiole shape in figs or aggressive behavior in ants. Nevertheless, the obligate mutualism implies full sympatry between the two partners, preventing within species comparisons of traits variations in absence of the mutualistic partners. A striking example of non-obligate mutualism driving trait convergence between sympatric species is Müllerian mimicry, whereby individuals from different chemically-defended species display similar warning color patterns (Müller 1879). This evolutionary convergence is driven by the predators learning the association between warning coloration and distastefulness, resulting in mutualistic relationships between sympatric species (Sherratt 2008). Mimetic interactions strongly depend on the local communities of defended species, resulting in spatial variations in mutualistic interactions and geographical variations in warning coloration (Sherratt 2006). In closely-related mimetic species, sharing a common coloration in sympatry may nevertheless lead to reproductive interference, because warning coloration is often used as a mating cue (Jiggins *et al*. 2001), and mimicry between species may thus enhance heterospecific sexual interactions. The costs generated by reproductive interference may thus limit convergence in warning colorations among closely-related species. The evolution of warning coloration could then be influenced by the relative abundance of sympatric species, modulating the positive effect of mimicry and the negative effect of reproductive interference on convergence in warning coloration.

Here we focus on three closely-related Neotropical butterfly species, namely *Morpho helenor* (Cramer, 1776), *M. achilles* (Linnaeus, 1758), and *M. deidamia* (Hübner, 1819), that exhibit substantial variation in dorsal wing colour patterns both within and among species, and whose large distribution ranges comprise situations of allopatry and sympatry (Blandin 2007). Bright coloration of wings is often associated with protection against predators either through chemical defenses or resemblance to chemically-defended species (Briolat *et al*. 2018). Although chemical defenses have not been reported in the three *Morpho* species studied, their iridescent blue band against a black background of the dorsal side of the wings is very conspicuous and strikingly contrasts with the cryptic colour pattern displayed by the ventral side (Debat *et al*. 2018). The flap-gliding flight behavior observed in these species (Le Roy *et al*. 2019) generates alternative phases of (1) flashes of light when wings are open and (2) vanishing when wings are closed. Associated with fast and erratic flight trajectories, these contrasted dorsal and ventral wing colour patterns make these butterflies difficult to locate and catch by humans and birds (Pinheiro 1996; Murali 2018). Wing colour patterns may thus induce predator confusion, enhancing escape capacities (Pinheiro *et al*. 2016). It was also suggested that such colour pattern might in turn act as a signal of such high escape capacities, further limiting predation attempts: Pinheiro & Campos (2019) observed that *M. achilles* and *M. helenor* butterflies were indeed frequently sight-rejected by wild jacamars. Since wild jacamars are important butterfly predators occurring in rainforests where these *Morpho* species are found, we can hypothesize that these predators could already have associated their wing patterns with high escape abilities. Behavioural experiments in controlled conditions have also shown empirically that predators learn to refrain their attacks towards preys displaying conspicuous coloration similar to that of previously missed prey (Gibson 1980). As in chemical-defense, escape ability can thus be associated with coloration by predators. Mimicking such a signal displayed by a prey with high escape abilities could thus provide protection against predators. Individuals sharing a locally abundant coloration associated with high escape capacities might thus benefit from increased protection against predators in the wild, favouring the persistence of similar colour pattern in sympatric species, as observed in chemically protected species (Müller 1879). Such ‘escape mimicry’ has been hypothesized in *Morpho* but never formally tested (Pinheiro *et al*. 2016). The three *Morpho* species studied here display geographic variation in colour patterns within species and their ranges largely overlap, *M. helenor* showing a more expended range in central America as compared to the other two species (Blandin & Purser 2013). *M. helenor* and *M. achilles* are sister species whereas *M. deidamia* belongs to a more divergent clade (Chazot *et al*. 2016) but nevertheless displays similar variation in dorsal color pattern. This situation allows comparing intra and inter-specific variations and testing the effect of sympatry on the evolution of traits across those closely related species. When these species occur in sympatry, they share the same micro-habitat (DeVries *et al*. 2010) and thus probably face similar communities of predators, enabling the evolution of mutualistic species interaction. In this study we test whether colour pattern variations across geographical areas observed in these three species are consistent with the hypothesis that local selection exerted by these shared predators promotes the evolution of convergent wing colour patterns, possibly *via* escape mimicry.

Based on the collection of *Morpho* held at the National Museum of Natural History in Paris (France), we finely quantified dorsal wing colour pattern of 723 specimens sampled throughout the whole distribution of the three species. We then specifically test the effect of sympatry on (1) colour pattern similarity between species pairs, by comparing phenotypic distance between species within and between localities (2) colour pattern variation in *M. helenor*, by comparing *M. helenor* populations where the other two species co-occur *vs*. populations where the other two species are absent.

## Material and Methods

### Sampling zones and specimens

The genus *Morpho* is distributed through three biogeographical regions: the Atlantic Forest region, the *cis*-Andean region (the Amazon and Orinoco basins, and the Guiana shield), and the *tran*s-Andean region (central and western Colombia, western Ecuador and north-western Peru, Panama Isthmus and Central America) (Blandin 2007; Blandin & Purser 2013). *M. helenor* is the only species covering the whole range of the genus, from northern Argentina to a large part of Mexico. It is also the most diversified species, with more than 40 described subspecies (Blandin 2007). *M. achilles* and *M. deidamia* exist only in the *cis*-Andean region, where they both are sympatric with *M. helenor* at the understory level in rainforests, from sea level (in the Orinoco delta) to more than 1000 m.a.s.l. in Andean slopes. However, *M. helenor* exists alone in dryer contexts, notably in the middle Marañon valley (Peru) and in eastern Venezuela.

We used the collections of National Natural History Museum of Paris to study the variation of colour pattern in these three species throughout their geographical range. These three species are generally locally abundant, and the Museum collection puts together numerous specimens collected at different times by different collectors. The possible impact of collectors’ bias towards rare wing patterns is thus probably limited in our sample.

Sympatry was defined as the co-occurrence of the three species within a *sampling zone*. We then defined *sampling zones* based on the geographical distribution of 17 subspecies of *M. helenor* (fig. 1): because subspecies of *M. helenor* are mostly defined based on colour pattern variation, sampling zones thus correspond to geographic regions where *M. helenor* butterflies display a similar colour pattern. In the absence of spatial heterogeneity in natural selection acting on colour pattern, we do not expect any increase in resemblance with the other two species within such sampling zones. Contrastingly, if escape mimicry promotes local convergence in colour pattern, the geographic variation observed in *M. helenor* is predicted to be mirrored by parallel variation in the other two species. Because in the *cis*-Andean region, *M. helenor helenor, M. h. theodorus*, and *M. h. coelestis* have very large geographical ranges (Blandin 2007), we further split these zones respectively into 2, 4, and 2 sampling zones. The total sample was composed of 22 sampling zones. We selected 723 specimens of *M. helenor* (n = 413), *M. achilles* (n = 156) and *M. deidamia* (*n* = 154), focusing on intact, well-conserved individuals (see supplementary table 1 for detailed numbers, and comments on the sampling zones). We used both males (*n* = 524) and females (*n* = 199), although females are less numerous in Museum collection because they are less frequently caught.

**Figure 1:**
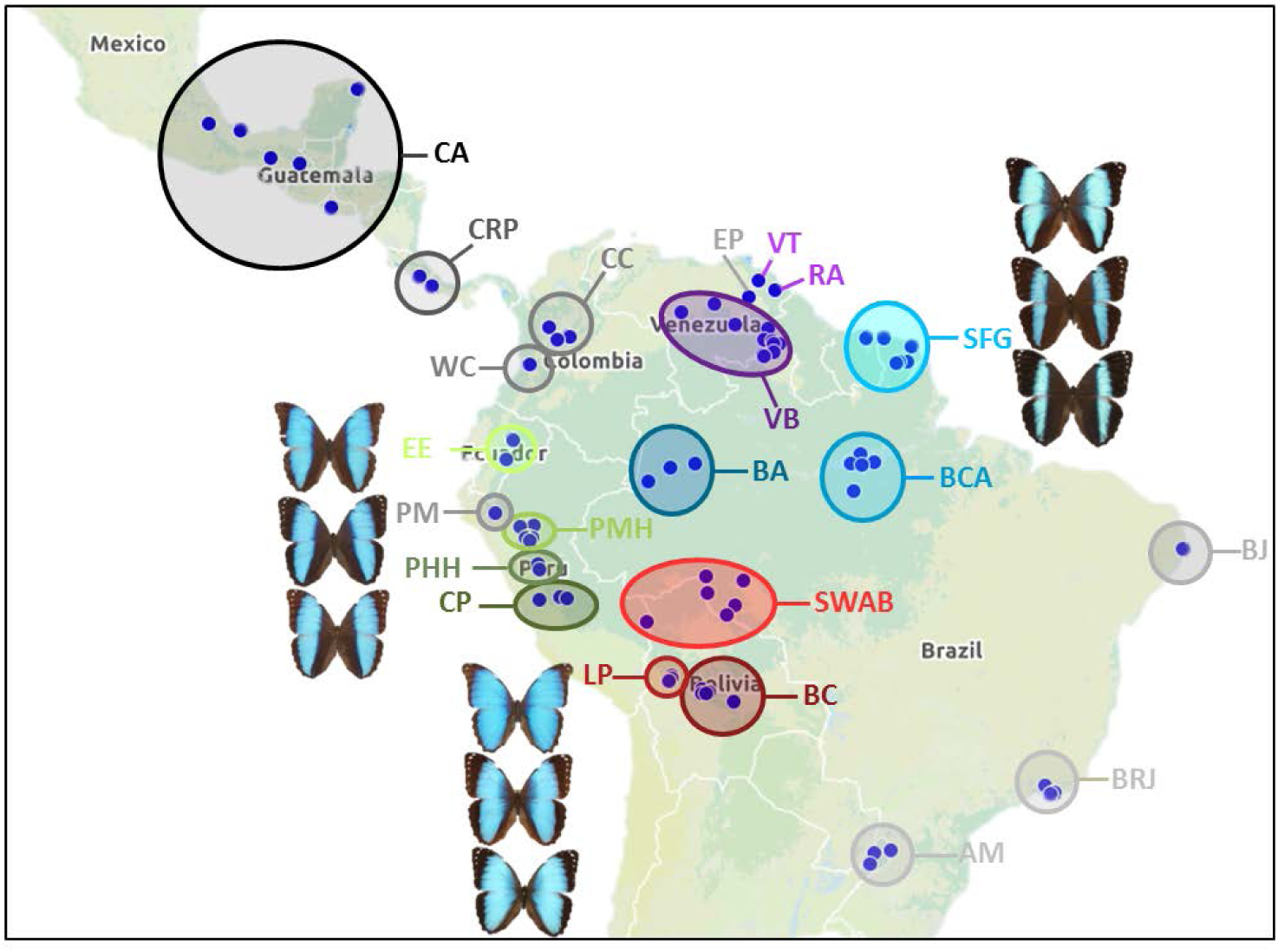
Geographic location of specimens, showing the 22 defined sampling zones. (see supplementary table 1 for detailed numbers of specimens per sampling zone). Sampling zones where the three species (*M. achilles, M. deidamia* and *M. helenor*) co-occur, and share the same micro-habitat, are shown with different colours whereas sampling zones where only *M. helenor* occurs are shown with grey levels. Examples of butterflies from the three species – *M. deidamia* (top) *M. helenor* (middle) and *M. achilles* (bottom) – sampled in the middle Huallaga in Peru (top left triplet), Bolivia (bottom left triplet) and French Guiana (top right triplet) are shown.

### Quantifying colour pattern variation

Pictures of collection specimens were taken in controlled standard white light conditions. The four wings were first manually separated using Adobe Photoshop Element. Wing images were then analysed following the Colour Pattern Modelling approach (Le Poul *et al*. 2014) implemented in Matlab. This method allows precise comparison of colour pattern while accounting for wing shape and venation that might differ between species. It has been shown to be especially relevant to quantify similarity and differences in color pattern within and across species (Le Poul *et al*. 2014; Huber *et al*. 2015; McClure *et al*. 2019). Briefly, the algorithm detects the four wings on the white background and segments the colour pattern in different categories based on pixel densities of the RGB values. The number of colours is then set manually: here we chose to consider three colours, namely black, blue and white. Some individuals (as for instance *M. deidamia* samples from French Guiana), display a gradient of blue (see sup. Fig. 1) that is often detected as a different colour category by CPM. Dark blue was nevertheless treated as blue in our analyses. This is probably a conservative assumption regarding convergence in colour patterns, because the dark blue area of *M. deidamia* has a similar location to the basal black area in *M. helenor* and *M. achilles*, and look very dark from far-distance. After segmentation, wings were aligned by adjusting translation, rotation and scale in order to maximize similarity, allowing the colour value for each pixel of the wings to be compared.

### Testing phenotypic convergence between species when living in sympatry

PCA based on colour values for each pixel on the four wings was then performed using the software R (R Core Development Team 2005), creating a morphospace where individuals located close to each other have a similar colour pattern. A MANOVA on pixel values observed on the 723 individuals from the three species was performed to test the effects of species, sex and sampling zone, as well as the interactions between all these variables. Under the hypothesis of convergent evolution of wing colour patterns in sympatric species, the levels of intra-specific phenotypic variation within sampling zones should be similar in the three sympatric species: this is expected as a result of positive frequency-dependent selection favouring similar phenotypes in the three sympatric species, but also because for each species, the protection gained by each phenotype depends on the range of phenotypes encountered by predators, which directly depends on the variation within the other two mimetic species. We used the trace of the within species covariance matrix based on the PCA axes as an estimate of the level of phenotypic variation within species within each sampling zone. We then computed the Pearson correlations between phenotypic variances observed in pairs of species across the 13 sampling zones where the three species co-occur. Significant correlations would be consistent with the hypothesis of convergent evolution of wing colour pattern in sympatry.

The phenotypic resemblance between species in sympatry was estimated by computing the average Euclidian distance between species within and across sampling zones in the PCA space (using the 15 first PCA axes, each explaining more than 0.5% of the total variance). To test whether species were more similar within a sampling zone than expected by chance, we generated a null distribution of distances between pairs of species by permuting the sampling zones within each species independently, so that the sympatry/allopatry relationships between inter-specific pairs of populations were randomized. We performed 10,000 simulations and computed within each simulation the average phenotypic distance between the three species pairs within sampling zones. This allowed generating for each pair of species, an estimated distribution of interspecific phenotypic distances within sampling zone under the null model assuming independent geographic variation in colour pattern within each species. We then assessed significance by counting the proportion of inter-specific distances under this null model that exhibited a lower value than the observed Euclidian distances between species within sampling zone. The observed interspecific distance was considered significantly smaller within sampling zone when the observed value was lower than 95% of the resampled values.

### Testing the effect of sympatry on wing pattern in M. helenor

Because *M. helenor* distribution range exceeds that of the two other species, it can be found in isolation in a significant proportion of its distribution (notably in Central America and in the East Coast of Brazil – see fig. 1). This situation allowed us to test the effect of sympatry on *M. helenor* colour pattern. We thus contrasted sampling zones where it co-occurs with the other two species (*i*.*e*. sympatric populations) with sampling zones where it occurs alone (*i*.*e*. allopatric populations). This effect of sympatry *vs*. allopatry on colour pattern was first explored in a PCA and then tested using a MANOVA, controlling for the effect of sex and sampling zone.

Under the convergence hypothesis, we also predicted that the phenotypic variance within *M. helenor* populations should be reduced when they are in sympatry with the two other species, because of the constraining effects of the selection imposed by other two species. We thus compared the level of phenotypic variation in *M. helenor* between sympatric and allopatric sampling zones. We thus computed the phenotypic variance within each *M. helenor* population using the trace of the covariance matrix of PCA coordinates and test whether the variances observed in the 13 sympatric groups of populations had smaller values than those observed in the 9 allopatric groups of populations using a simple *t*-test.

## Results

### Geographic variation of wing colour pattern across the three species

The PCA based on colour variation in wing pixels shows that individuals sampled within a sampling zone tend to display a similar wing colour pattern (fig. 2). This effect was confirmed by the MANOVA: We detected a strong and significant effect of sampling zone (*Pillai* = 13.21, *F* = 2.87, *df* = 81, *P* < 0.001), species (*Pillai* = 1.27, *F* = 19.94, *df*=2, *P* < 0.001) and sex (*Pillai* = 0.80, *F* = 46.51, *df*=1, *P* < 0.001), as well as significant interactions between species and sex (*Pillai* = 0.67, *F* = 5.84, *df*=2, *P* < 0.001), species and sampling zone (*Pillai* = 4.74, *F* = 1.86, *df*=36, *P* < 0.001), as well as sex and sampling zone (*Pillai* = 5.72, *F* = 2.06, *df* = 40, *P* < 0.001). The observed geographic variation is thus consistent with a greater resemblance between individuals within a sampling zone as compared to individuals sampled in different sampling zones, modulated by some sexual differences and variations between species.

**Figure 2:**
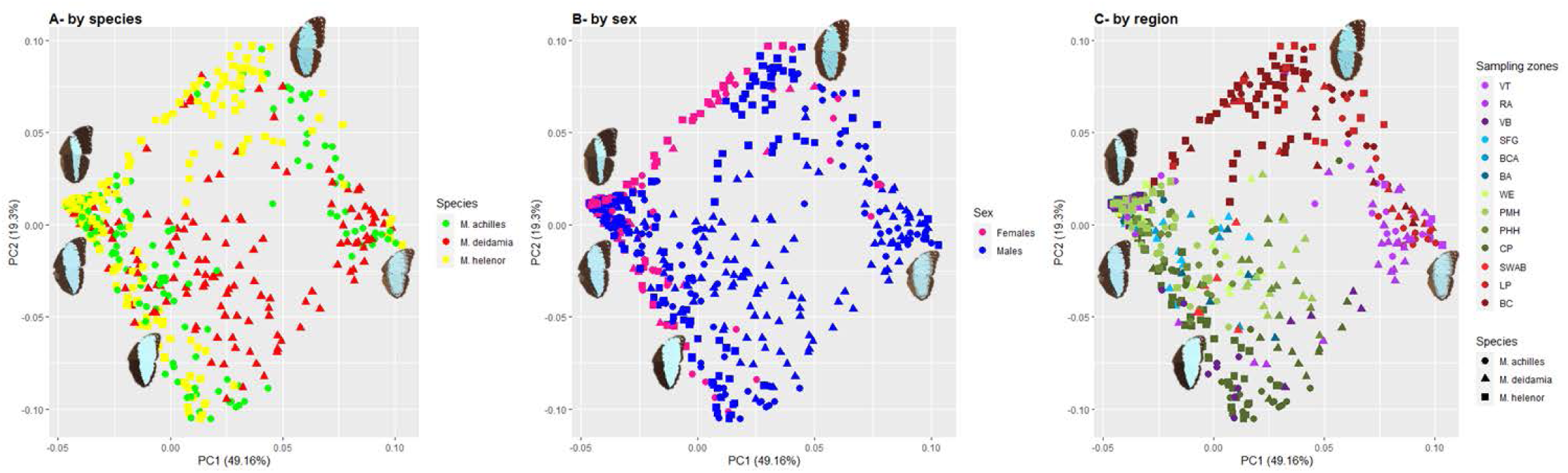
Colour pattern variations among individuals living in sympatry, from different species (A), sexes (B) and sampling zones (C), captured by the PCA based on pixels colour variations analysed by CPM. Only sampling zones where the three species live in sympatry are represented here (n = 557 individuals). To explicit phenotypic variations, male specimens of *M. achilles* sampled in different sampling zones are shown closed to their location on the morphospace. Symbols differ among species, with dots for *M. achilles* individuals, triangles for *M. deidamia* and squares for *M. helenor*. The colours of symbols differ among species (left plot A), sex (central plot B) and sampling areas (right plot, C). Note the colours of sampling areas on plot C match the colour code used on the geographic map (fig.1).

In the 13 localities where the three species co-occur, the phenotypic variance within species was highly correlated between *M. helenor* and *M. achilles* (Pearson correlation: *cor* = 0.83, *P* = 0.0015) and *M. helenor* and *M. deidamia* (Pearson correlation: *cor* = 0.83, *P* = 0.0012), and moderately correlated between *M. deidamia* and *M. achilles* (Pearson correlation: *cor* = 0.66, *P* = 0.028). This suggests that within sampling zones, the level of within species phenotypic variation is similar across species. This could stem from a sample bias among sampling zones, but is also consistent with the convergence hypothesis, whereby phenotypic variation within a species indirectly impacts the phenotypic variation in sympatric species by modifying the selective pressure.

### Convergence of colour patterns between species within sampling zones

To investigate the hypothesis of local convergence among the three species more directly, we then specifically tested whether the mean Euclidian distance between species within sampling zone was lower than expected when assuming an independent geographic differentiation in colour pattern within each species. Considering the whole dataset (both sexes together and including all sampling zones), the average phenotypic distances between each pair of species within sampling zone were all significantly smaller than under the simulated null distribution (fig. 4). These tests were also significant when excluding the sampling zones where *M. helenor* occurs alone (see supplementary figure 1), and when carried out separately on males and females (see supplementary figures 2 and 3 respectively), confirming the significant convergence between species in sympatry. The observed inter-specific phenotypic distances in sympatry were lower between *M. helenor* and *M. achilles* than between the other two pairs of species, consistent with their closer phylogenetic distance (fig. 4). Despite this strong signal of convergence of colour pattern in sympatry, some sampling zones departed from this general trend, notably the zones situated in Venezuela (VT, VB and RA) and Bolivia (LP).

**Figure 3:**
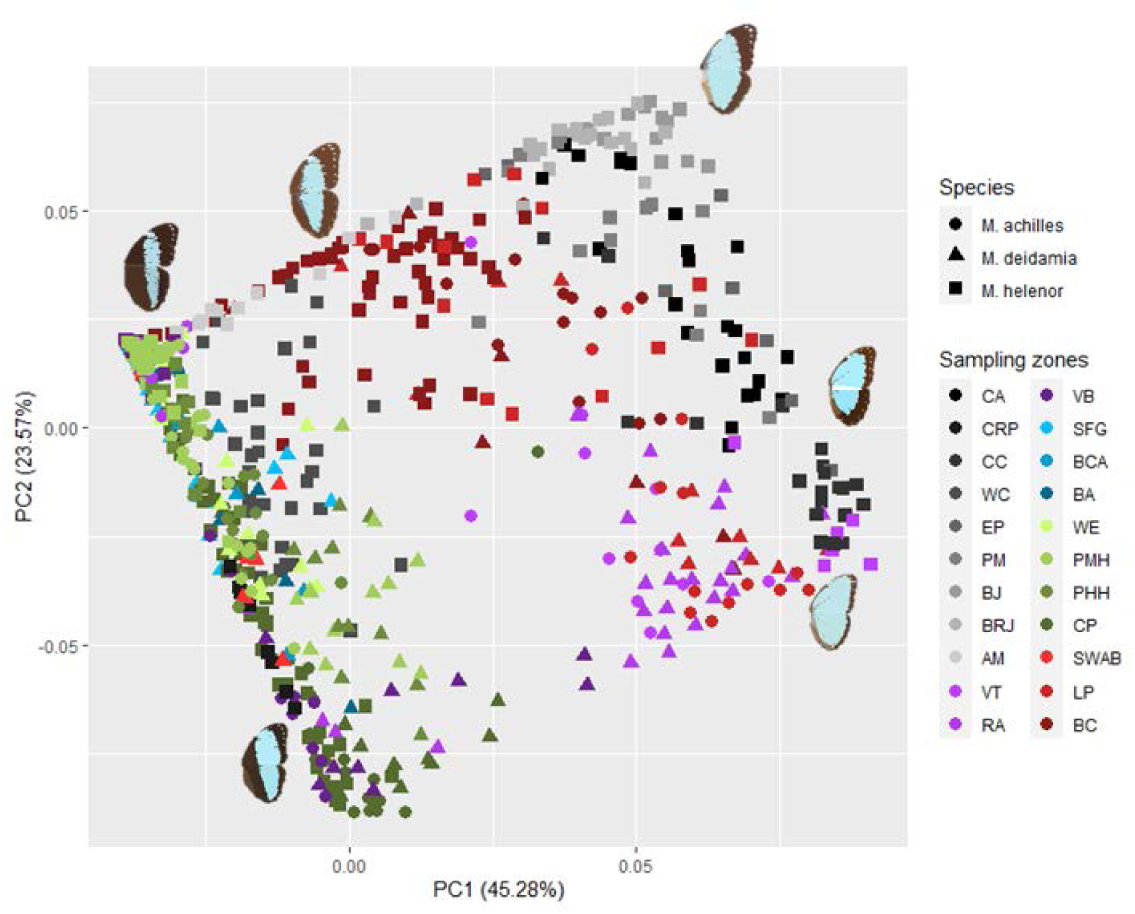
Colour pattern variations among individuals in sampling zones where *M. helenor* is in sympatry with the other two *Morpho* species (colored symbols) and in sampling zones where *M. helenor* does not co-occur with the other two species (grey-scale symbols), from the three different species represented by the first two axes of the PCA based on pixels colour variations analysed by CPM (*n* = 723). To explicit phenotypic variations, male specimens of *M. helenor* sampled in different sampling zones are shown closed to their location on the morphospace. Symbols differ among species, with dots for *M. achilles* individuals, triangles for *M. deidamia* and squares for *M. helenor*. Note the colours of sampling zones match the colour code used on the geographic map (fig.1).

**Figure 4:**
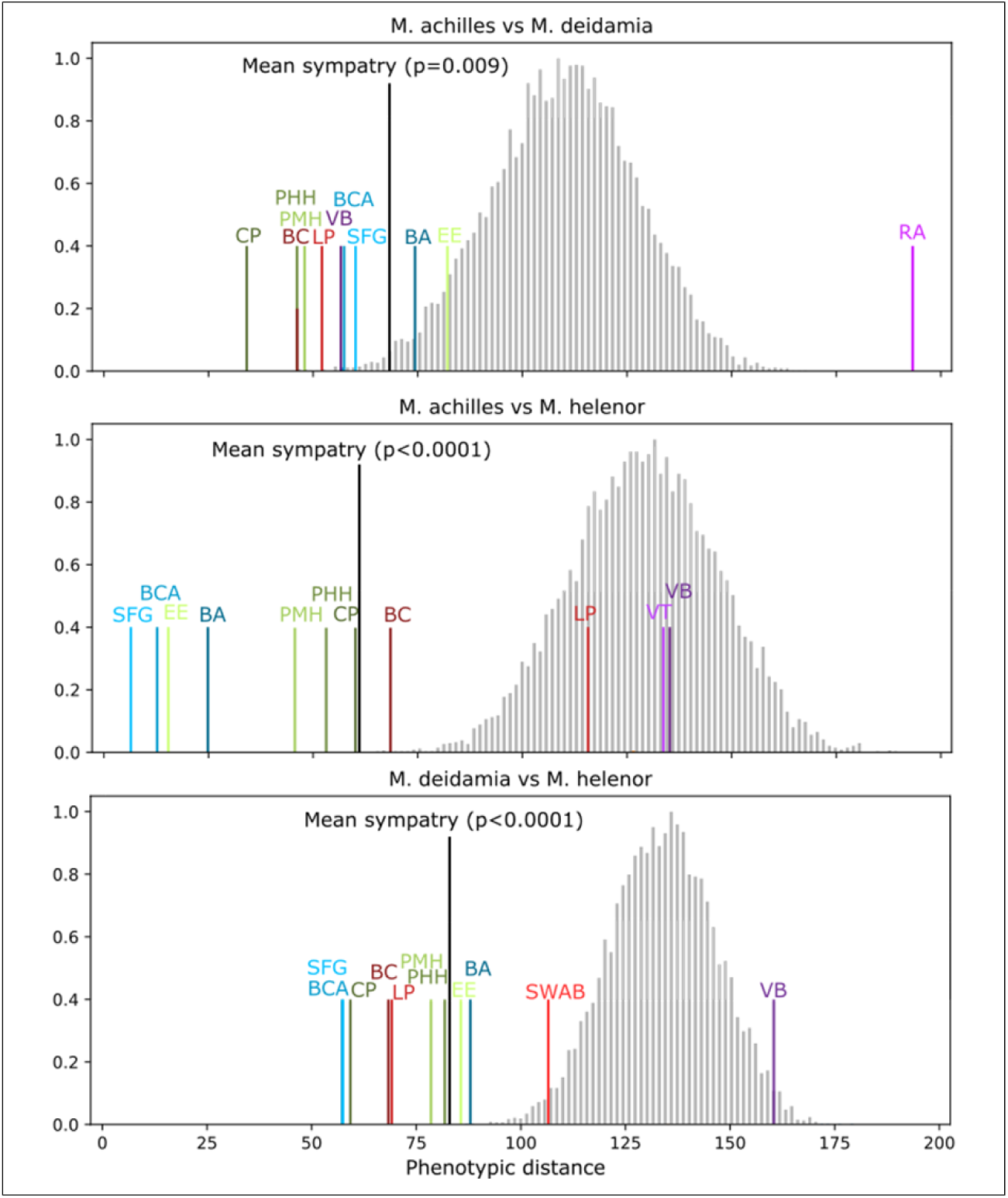
Phenotypic distances between species in sampling zones where they are found in sympatry (colored bars) and predicted distribution obtained using 10,000 bootstraps, randomly reallocating the different sampling zones within each species (grey bars). The black bar shows the mean phenotypic distance observed between pairs of species in the sympatric sampling zones; top plot: distances between *M. achilles* and *M. deidamia*, middle plot: distances between *M. helenor* and *M. achilles*, and bottom plot: distances between *M. deidamia* and *M. helenor*. The *p*-value is based on the number of simulations where the phenotypic distance between species is higher than the mean value of inter-specific distances observed in sympatry. Note the colours and codes of sampling zones match the colour code used on the geographic map (fig.1).

### Effect of sympatry on colour pattern variation in M. helenor

We then compared the colour pattern of *M. helenor* from sampling zones where it co-occurs with *M. deidamia* and *M. achilles* to that of sampling zones where it occurs alone, using a PCA on sympatric and allopatric individuals of the three species (*n* = 723). Interestingly, some allopatric populations of *M. helenor* (in particular populations located in the Atlantic coast of Brazil, *i*.*e*. BJ and BRJ) show wing patterns that differ from individuals living in sympatry in other geographical areas, as highlighted by their distributions in the wing colour pattern morphospace (fig.3). Using a MANOVA on colour pattern variations in *M. helenor* only (*n* = 413), a significant effect of sympatry was detected (*Pillai* = 0.95, *F* = 155.77, *df* = 1, *P* < 0.001), controlling for the effect of sex (*Pillai* = 0.71, *F* = 19.41, *df* = 1, *P* < 0.001) and sampling zone (*Pillai* = 7.81, *F* = 5.66, *df* = 19, *P* < 0.001).

The levels of phenotypic variation in *M. helenor* were also slightly higher in allopatric populations (*mean variance* = 4766) as compared to sympatric populations (*mean variance* = 2571) (*t* = 1,997, *df* = 18.96, *P* = 0.06). Although neutral divergence among these geographically distant localities might contribute to the observed effects, these comparisons between sympatry and allopatric populations of *M. helenor* are consistent with a substantial effect of species interactions on the evolution of wing colour pattern in *M. helenor*.

## Discussion

### Estimating phenotypic variation based on Museum collections

Our study of phenotypic variation within and among species was enabled by the rich *Morpho* collection held in the Museum of Natural History in Paris, containing large number of specimens collected throughout the whole geographic distribution of the three species studied here. Nevertheless, estimations of phenotypic variation based on museum collections can be biased: (1) females are indeed generally under-sampled, because they are less frequently encountered in the wild; (2) phenotypic variation can be overestimated, because collectors tend to prefer specimens with unusual colour patterns. Concerning the first bias, the sexual dimorphism in colour pattern is very limited in the three studied species (fig. 2B) and the signal of convergence observed was similar when considering males and females separately, suggesting that the convergence observed is likely to occur similarly in both sexes. Concerning the second bias, an overestimation of phenotypic variation is likely to decrease the signal of convergence; therefore our approach based on collection specimens is probably conservative relatively to phenotypic convergence.

### Local similarity of colour pattern between species

By precisely quantifying colour pattern variation across a large sample of *Morpho* butterflies, we detected a significantly increased resemblance between individuals from different species living in sympatry as compared to allopatry. This convergence is stronger between the two sister species *M. helenor* and *M. achilles* than for the more distantly related species *M. deidamia*, probably because the larger phylogenetic distance might involve stronger developmental constraints. This convergence trend is confirmed for a majority of sampling zones located in the Amazonian rainforest, resulting in similar ecological conditions encountered by *Morpho* butterflies in these different geographic areas. Despite this similarity in ecological conditions, wing colour patterns displayed by butterflies from different species are more similar within localities as compared to across localities, suggesting that convergent evolution of colour patterns might be promoted throughout the three species.

In Bolivian and Venezuelan sampling zones (LP, RA, VT, VB, see fig. 1), a large diversity of colour pattern was observed within species. Although such a high level of intraspecific variation is found in the three species (fig.2C), it is likely responsible for the non-significance of our test comparing inter-specific phenotypic distance within and among sampling zones (fig. 4). However, the colour patterns displayed by all three species are rather similar, and quite different from those observed in other geographic regions (fig. 2C). The important variation within each species in these populations might thus still be consistent with the convergence hypothesis.

Overall, (1) the greater resemblance between the three species within localities in the largest part of their common range, together with (2) the divergence in colour pattern displayed by *M. helenor* butterflies in localities where the other two species do not occur, point at a role of species interactions in the evolution of dorsal wing pattern in the three species.

### Local convergence: escape mimicry or other shared local selective pressure?

How can we account for the similar variation among populations of the three *Morpho* species? One hypothesis would be that similar environments result in shared selective pressures acting locally on the three species, leading to the observed similarity of colour patterns, independently from interactions occurring in sympatry. What local selective pressures might be involved is however unclear. It has been shown that the evolution of warning coloration can be influenced by selective forces independent from mimetic interactions. For instance, the light environment may modify the conspicuousness of colour patterns (Rojas *et al*. 2014) so that variations in light environment in different localities may select for different wing colour patterns. However, the three *Morpho* species studied here mostly fly in the understory, regardless of the geographic regions, suggesting that the different populations may be evolving in similar light environment. Another hypothesis could stem from the role of melanin in thermoregulation (e.g. in Colias butterflies, Ellers & Boggs 2004) that may result in contrasted selective pressures in different geographic regions. The variation in dorsal pattern observed among *Morpho* populations indeed mostly affects the proportion of melanic patches on the wing, the populations of Surinam and French Guiana being the darkest (SFG locality on fig. 1). Variation in melanic surface on butterfly wing has been related to adaptation to cold environments in butterflies (e.g. in *Parnassius phoebus*, Guppy 1986). However, populations of *M. helenor, M. achilles*, and *M. deidamia* with very reduced black areas occur at sea level in the Orinoco delta, as well as around 700-800 m.a.s.l. in Bolivian valleys. Moreover, the darkest specimens occur at low altitudes in French Guiana, while populations with wider blue bands occur in some Peruvian valleys at more than 1000 m.a.s.l. The extension of black areas in these *Morpho* species therefore does not seem to occur in colder environments, making inconsistent the hypothesis of an effect of adaptation to temperature on black colour pattern evolution.

Although we cannot rule out that an unidentified local selection might promote the evolution of a similar colour pattern in the three species, the observed repeated local convergences are also consistent with the escape mimicry hypothesis. In Müllerian mimetic species such as the butterfly species *Heliconius melpomene* and *H. erato*, multiple geographic races with striking colour pattern variations are maintained within species, with strong resemblance to races from the other species (Jiggins 2017). These multiple locally convergent colour patterns are maintained by positive frequency dependent selection due to increased protection of mimetic colour patterns, reinforced by sexual preferences toward locally mimetic mates (Merrill *et al*. 2012). The geographic variations of these mimetic coloration then mainly stem from stochastic processes, because the colour pattern may not necessarily provide a selective advantage *per se*, but can be favored once it becomes frequent within a given locality where predators learn to avoid it (Mallet 2010). These evolutionary forces documented to drive variations in mimetic coloration in defended species might explain the multiple convergences observed in the three *Morpho* species throughout their geographical range. There might be no specific selective advantage related to the extent of the black bands on the wings, which might vary randomly across localities. The local convergence might in turn be driven by local positive frequency-dependent selection favouring the most common phenotype – the most avoided by predators. Our results are thus consistent with the escape mimicry hypothesis, whereby, similarly to chemical defenses, the high escape capacities of *Morpho* butterflies would promote local convergence of colour patterns.

### Convergent wing patterns in closely-related species

Because the three *Morpho* species studied here are closely related, their local resemblance is probably facilitated by some common developmental bases of colour pattern variation and could also be favored by local gene flow, explaining why convergence is weaker in the more distantly related species *M. deidamia*. The genetic basis of colour pattern variation in each of the three species needs to be identified to infer the level of independent phenotypic evolution in these three species. But interestingly, the increase in the extent of the black bands, observed for instance in butterflies from French Guiana and Surinam (SFG, fig. 1) seems to result from a different developmental process in *M. deidamia*, as compared to *M. helenor* and *M. achilles*: in *M. deidamia* the reduction of the blue band partly results from the development of a dark blue band whereas the similar reduction of the blue band of *M. helenor* and *M. achilles* stems from an extension of the area where black melanic scales are produced (see sup. fig. 1). This suggests that the apparent phenotypic similarity of dorsal colour pattern might arise at least partly from different developmental processes, as expected from convergent evolution.

If escape mimicry plays a role in the observed similarity between species within some localities, the evolution of colour pattern in these *Morpho* species would then strongly depend on the history of their sympatry in different geographic areas. When the different species are ancestrally allopatric, the evolution of colour pattern would then follow an advergence scenario, where the evolution of wing patterns in recently settled species would be strongly constrained by the colour pattern exhibited by an already abundant species. Alternatively, parallel geographic diversification in colour patterns might have occurred if these three species have been sympatric throughout their evolution and expanded their geographical range simultaneously. The history of wing pattern diversification in the three species thus strongly depends on their biogeographic history and in particular, on the history of speciation in this clade.

Because wing colour patterns are frequently used in butterflies as a mating cue, the high resemblance between closely-related species might lead to costly reproductive interference (i.e. interspecific courtship or even mating). Such costs of reproductive interference are thus predicted to limit the convergence triggered by mimicry: divergence in colour pattern would thus be favoured during speciation or in case of secondary contacts (Lukhtanov *et al*. 2005). This balance between mimicry and reproductive interference might explain the limited convergence between the three *Morpho* observed in in this study in some localities.

## Conclusions

By extensively studying wing colour patterns in three closely-related *Morpho* butterflies species throughout their geographic range, we detected significant resemblance among species within some localities and parallel variations across localities. Those results are in line with the escape mimicry hypothesis, which assumes that the similar high escape capacities of these three species may promote mimicry in their dorsal colour pattern. Besides providing evidence for convergence in colour patterns in *Morpho* in line with the escape mimicry hypothesis, our study more generally highlights the effect of sympatry on phenotypic evolution across species, stressing the need to jointly consider intra and interspecific variations to understand phenotypic evolution in sympatric species.

## Acknowledgments

The authors would like to thank Stephen Panara, Nicolas Zimmerman, Charlotte Ballin for editing pictures of *Morpho*. They are also grateful to Marianne Elias, and Nicolas Chazot for scientific discussions on trait diversification, as well as Nicolas Navarro and Sylvain Gerber for thoughtful discussions on estimations of phenotypic distances. This works was funded by the Emergence program of Paris city council to VL. All authors declare having no conflict of interest.

